# FPGA-based scanner and SerialEM server for 4D-STEM Electron Tomography

**DOI:** 10.64898/2026.06.26.734744

**Authors:** Shahar Seifer, Michael Elbaum

## Abstract

Four-dimensional scanning transmission electron microscopy (4D-STEM) enables the acquisition of diffraction patterns at every probe position in a dense array. For imaging applications this approach offers significant benefits in terms of spatial resolution and contrast enhancement. In this work, we present the development of a synchronous scan generator integrated with SerialEM software to enable automation of complex experimental protocols such as tomography. The proposed hardware functions as an interface between SerialEM, the scan controls of the microscope, a fast annular dark-field detector, and a synchronized trigger for a pixelated detector. Our previous implementation, named SavvyScan, relied on a dedicated computer equipped with a multichannel acquisition and signal-generation cards, as well as a separate microcontroller for synchronization. Here, we report a low-cost implementation based on a Red Pitaya board, utilizing direct programming of its embedded FPGA and Linux server components. We provide detailed instructions for system installation and operation, along with practical guidance for modifying the source code. System performance is validated through oscilloscope measurements and imaging of a replica grating sample. The utility of the approach is further demonstrated by generating a 3D electron tomogram of a cryogenic sample of mitochondria from a tilt series of shadow montage projections.

## Introduction

This work is part of our long-standing endeavor to advance the modality of scanning-TEM tomography for life-science cryogenic samples [1]. Cryo-scanning transmission electron tomography (cryo-STET) enables three-dimensional imaging of unstained, fully hydrated vitrified cells, in which flexible modes of STEM signal formation alleviate the thickness limitation inherent to conventional energy-filtered cryo-TEM. This provides access to larger cellular volumes with improved depth resolution, particularly at high tilt angles, while preserving native-state ultrastructure.

We consider the use of SerialEM software [2] to be central to the workflow, as described in [3]. The workflow supports low-electron-dose searching for features of interest, registration of target areas, and tracking. It also enables automated tilt-series acquisition for tomography by controlling sample tilting and tracking to stabilize the field of view. The software is compatible with electron microscopes from all major manufacturers and can connect to multiple camera servers, including STEM systems that generate images using a scanning probe. The Camera Setup interface in SerialEM allows users to define scan parameters such as dwell time, image size, and pixel sampling density. It further supports dynamic focusing in STEM by adjusting the microscope optics according to the beam position on a tilted sample. The hardware interface to a camera operates via a server model. Our SavvyScan system [4,5] is one such server, providing STEM images based on a selected detector. It incorporates an independent scan engine that connects to the *X* and *Y* channels of the microscope beam deflectors. The server integrates into SerialEM via a plugin, while operating as an independent process that stores additional datasets acquired simultaneously from multiple detectors. Accordingly, the output from an azimuthally segmented quadrant detector is used for integrated differential phase-contrast (iDPC) techniques [4,6], while the output from a pixelated detector enables 4D-STEM techniques [7,8].

Recent developments in 4D-STEM have shown clear practical advantages in high-resolution imaging and analysis [9–11]. By recording a full diffraction pattern at each probe position, the method captures more complete scattering information than conventional STEM, which can then be reconstructed in different modes from the same dataset. This enables improved contrast, particularly for weakly scattering and beam-sensitive samples. In addition, the use of phase-retrieval methods such as ptychography can enhance spatial resolution beyond that of conventional STEM techniques [12,13]. A variant of such approaches is our shadow montage technique that resolves layers in the sample by summation of diffraction patterns treated as cone-beam shadow projections [8]. A different analysis of similar raw data is used for the tilt-corrected bright field (tcBF) method based on parallax correction [10].

Furthermore, because the recorded signal retains angular scattering information, it can be analyzed quantitatively to provide material characteristics based on inferred elastic and inelastic scattering contributions [14]. Our aim has been to extend these capabilities to 3 dimensions by supporting tilt-series acquisition in 4D-STEM. Here we present a second-generation hardware platform that is at once flexible, inexpensive, and designed from the outset with SerialEM integration in mind. Based on a Red Pitaya development board, ironically named STEMlab, it is also compact and especially simple to implement.

## System Design

We briefly summarize the key features of the SavvyScan system described in our previous publication [4,5] and focus here on the design changes and the motivation for the new implementation. The system diagram is shown in Fig.1, in which the Red Pitaya board is connected via a local Ethernet network to a Windows workstation running SerialEM and to the 4D-STEM camera server. In our system this is the DECTRIS server unit for controlling an ARINA pixelated detector via HTML commands. Ethernet connection between SerialEM and the electron microscope is also required, optionally in a separate network. Two analog outputs of the Red Pitaya board are connected to the microscope scan controls, and a digital output serves for triggering the pixelated detector. As a scan engine, our software assumes a certain scan polarity according to the one available in Tecnai F20. In the Titan Krios, by contrast, the polarity is selected by a choice of socket.

**Figure 1.**
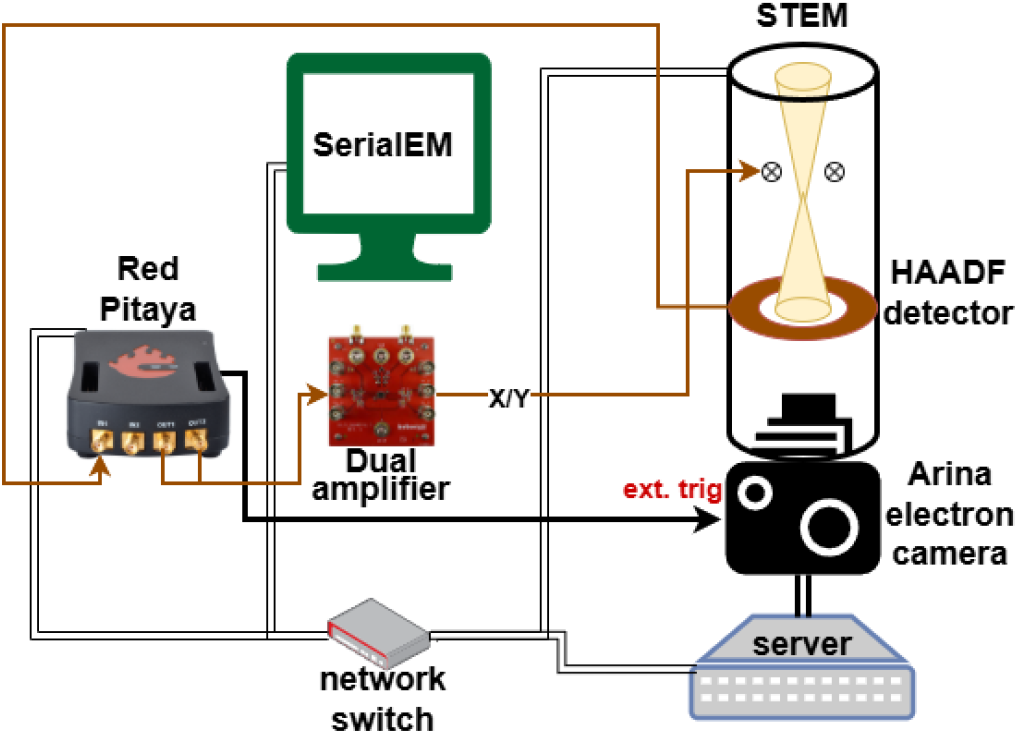
Scheme of the FPGA-based SavvyScan system and connections to the STEM microscope

SavvyScan is designed with flexibility as a core principle, most notably through programmable scan patterns. It was originally designed for simultaneous acquisition from multiple detectors, such as the Opal segmented diode (El Mul Technologies), and later adapted for 4D-STEM [5]. In the new FPGA implementation, we retain the same data structures as in the previous version. For a conventional raster, we define a scan pattern that begins outside the imaging area so that the scanning probe reaches steady velocity before entering the region of interest. Notably, the SavvyGate auxiliary provides a synchronized stream of trigger pulses only in the image area, to avoid the need to clean the data files later. The scan pattern is stored in on-board RAM and executed directly on the FPGA, providing for additional scan patterns such as zig-zag, spiral, sliding circles.

In contrast to the previous design, which supported eight analog-to-digital converter (ADC) channels, the current implementation uses a single ADC channel dedicated to the high-angle annular dark-field (HAADF) detector. The HAADF signal is selected as the source for image tracking and navigation. Diffraction images are acquired on a pixelated detector at selected scan positions defined by a gate pattern table. (The same feature might be used to control a fast electrostatic shutter for compressive sensing applications.) In the new register-transfer level implementation there is no lower bound for the scan rate, and a new feature allows variable trigger delay to support frozen beam acquisition at each scan point.

The new design simplifies user operation by optionally eliminating a separate user interface and reducing configuration elements. The goal is to provide a transparent workflow that can be operated in a manner similar to conventional electron tomography, without requiring extensive training in 4D-STEM techniques. All system operations are handled directly from SerialEM, and a dedicated 4D-STEM control is provided through SerialEM script functions.

The overall architecture is simplified through direct FPGA programming, which removes several constraints present in the earlier implementation. Technical details and instructions appear in the appendix. Over-sampling and averaging of the HAADF ADC signal within a single pixel dwell time, for noise reduction, is now performed directly on the FPGA. This significantly reduces the processing load on the server. Such an approach has shown improved signal to noise ratio in other scanning probe systems [15]. A limitation of the current implementation is the available on-board memory of 256 MB, which currently restricts the maximum scan size to 2048 × 2048 points. In practice this constraint is not limiting for typical applications, especially in light of the up-sampling made possible by defocused 4D-STEM methods such as ptychography, tcBF, and shadow montage.

### Operating Instructions

Connect and power up the Red Pitaya Board (RPB) and dual amplifier (the latter uses regulated ±12 V) according to the wiring scheme of Fig.1.

SSH into the RPB Linux server and type in the /root folder:

bash ./start

Wait for the server to start, then start SerialEM using the shortcut file in the Start_SerialEM folder.

Single images are acquired as normal in SerialEM using the Camera Setup and optionally the Low Dose controls. Take care that the HAADF detector is inserted and selected in the Camera Setup window, and adjust the HAADF contrast/brightness to proper range. The full digital range is supported by SerialEM in camera option “Divide 16-bit by 2”. Ensure that the external scan is enabled in order to bypass the internal scan generator. Typical settings include *Search* (quarter FOV, 256 × 256, 30 μs/px), *Trial* (full FOV, binned to 1024 × 1024, 6 μs/px), and *Focus* (quarter FOV, 512 × 512, 1s/frame). In *Record* the FOV is full, binned to 1024 × 1024 or 2048 × 2048. Note that the flyback time is included in the SerialEM display of the pixel dwell time. The ARINA camera reaches 8 μs frame speed only when operating in free-running mode. Using the external trigger to avoid acquisition during flyback and acceleration, we recommend a 14 μs trigger interval, which requires 20 μs dwell in the Camera Setup due to the extra flyback time.

Follow the instructions in SerialEM documentation for proper calibration of the stage and image shifts. If the calibration is already done in a different SerialEM camera the Savvyscan voltage range may be adapted by changing the related parameters in feeder.cpp (output_amplifiers_gain and parking_voltage) and compiling the code

bash ./compile

Ensure that the indicated voltage never exceeds the output range of the RPB, which is ±1 V, otherwise the output will be clipped.

A script file contains available actions that are not implemented in SerialEM but can be run from its script engine via CallFunction send_command <property>. The single-line calls are deactivated by comment characters (#) and include the properties listed in Table 1.

**Table 1.** Properties of the function send_command in SerialEM scripting

| Property | Description |
| --- | --- |
| ARINA: <filename> | Arm the ARINA camera and set a master filename. |
| StartTiltSeries: <number> | Start a tilt series, and if ARINA is armed keep this state for the specified number of tiltviews and write their counter in the hdf5 filename. |
| StopTiltSeries | End the tilt series prematurely. |
| SetThTime: <time in seconds> | Record 4D-STEM datasets for scan requests longer than this threshold. |
| SetTrigerDelayus: <time in μs> | Set a delay between the signal to the scan coils and a trigger to the pixelated detector (default is μs). |

Additional options related to communication with the ARINA are under development.

To start a tilt series in 4D-STEM, first use the ARINA and StartTiltSeries script commands with a unique filename. Then work with eucentric height calibration and tilt series acquisition available in SerialEM menu. Choose the tilt angle range and steps, and optionally set the dose-symmetric tilt order. (For maximum stability, particularly with a side-entry holder, select “Tracking by Trial scans only”). Then push the Run button and choose the mrc filename for storing the HAADF output. At the end, push Terminate to close the operation. Different protocols for single-particle analysis or micro-ED/3DED may be adopted straightforwardly by means of SerialEM scripting [16,17].

### Validation

The oscilloscope trace shown in Fig. 2 validates FPGA operation with precise synchronization between the analog output (AWG ch1) and the camera trigger signal (DIO3_P). The oscilloscope is triggered by the DIO0_P output, which is generated by the FPGA upon receiving a command cue in RAM to initiate the scan. The DIO3_P pulses are 3 μs wide and repeat for each pixel of the generated 1024 × 1024 HAADF image broadcast to SerialEM. Each burst of pulses corresponds to the acquisition of a single image line, where the final camera trigger ends immediately before the scan-x voltage returns to a position outside the field of view in preparation for the next scan line. This synchronization is maintained throughout the entire scan. At the conclusion of the scan, the beam is deflected to a parking position outside the field of view to prevent sample damage. An Arduino microcontroller was used to verify that exactly 1024^2^ trigger pulses are received.

**Figure 2.**
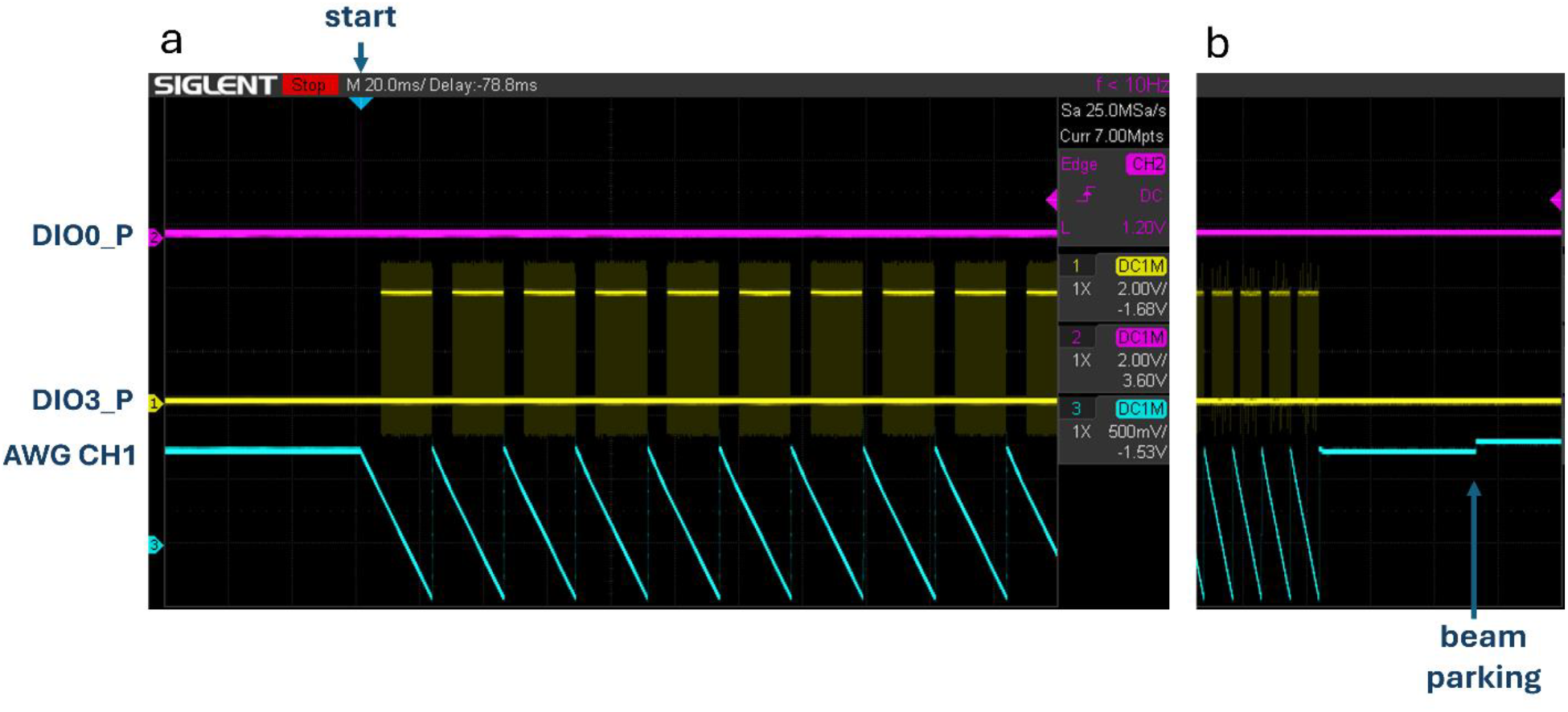
Signals on scope demonstrate synchronization during a scan: DIO0_P serves internally to start acquisition. DIO3_P is the trigger for the pixelated detector (ARINA) based on 1024 pulses in each line of the scan. AWG CH1 is driving the X scan coil of the microscope. Trace b shows the end of the scan in which the signals are synchronized and the beam parking state is engaged.

To test synchronization between the output and input channels one can immediately verify the correct imaging pattern by directly connecting one output channel to the input CH1. For accurate validation, we tested a standard replica grating sample (EMS) on a Tecnai-F20 S/TEM microscope (FEI, Inc) operating at 200 kV acceleration in NanoProbe mode with 30 μm C2 aperture and a HAADF detector (Fischione). The result shown in Fig.3 demonstrates straight grating lines that verify correct geometrical mapping and linearity of the dual amplifier (configured as an inverting operational amplifier with gain of -11), and the spatial calibration is consistent with the specified grating density of 2160 gratings mm^−1^. Clear contrast is observed between the carbon layer (gray) and the latex bead (white). Correct calibration is also validated by shifting the scan position and confirming agreement between the predicted and measured locations.

**Figure 3.**
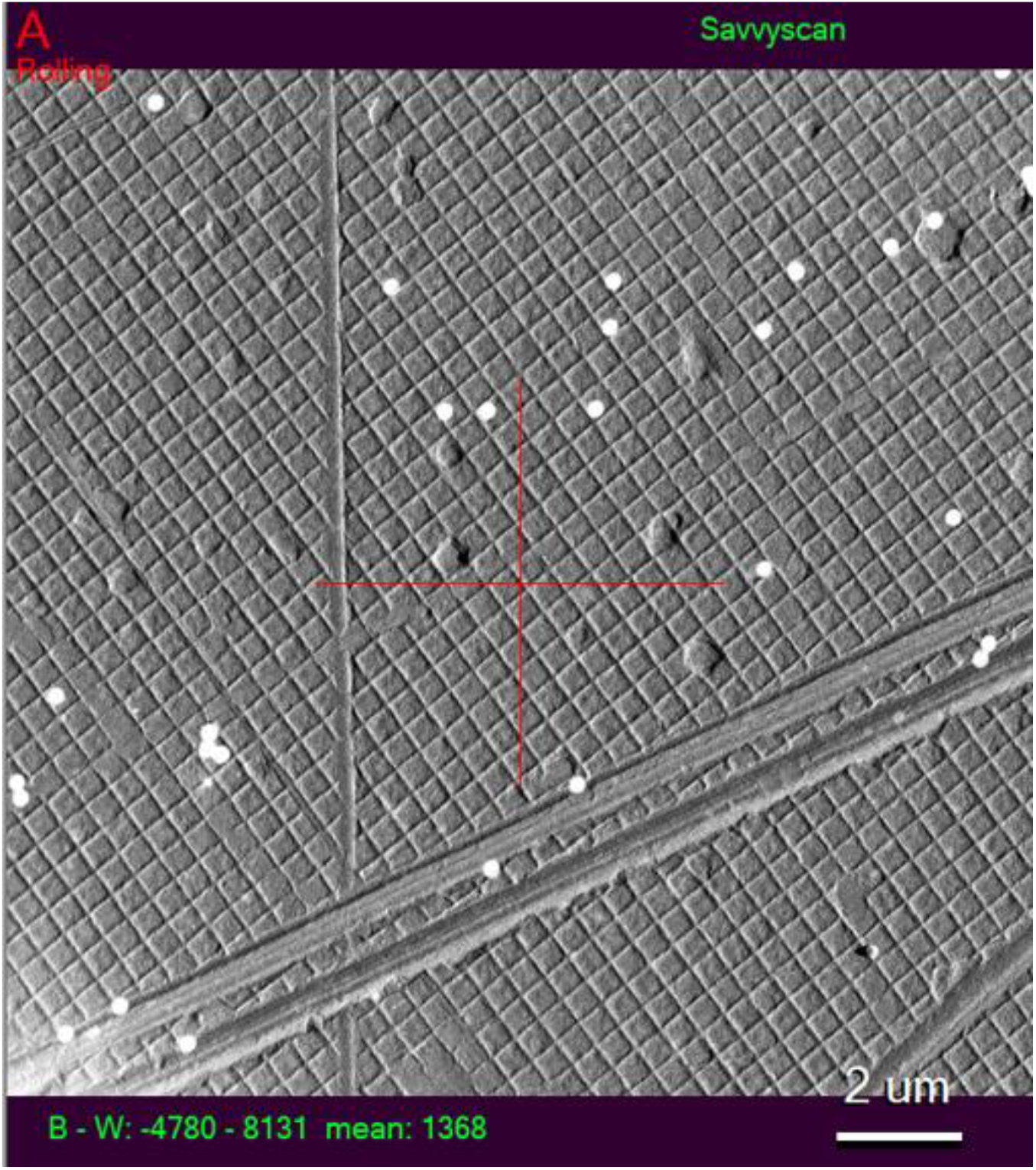
The output window in SerialEM after scanning of a standard replica grating. The image demonstrates geometric uniformity over the field of view, and a proper calibration based on the specified grating density of 2160 lines per mm.

At higher magnification, zooming into the center of the low-magnification image is demonstrated. The HAADF image shown in Fig. 4a was acquired using a 512 × 512 scan. Simultaneously, a 4D-STEM dataset was collected using the ARINA pixelated detector, comprising 512^2^ diffraction images of 96 × 96 pixels each. Applying the shadow montage technique to this dataset yields the image shown in Fig. 4b, which exhibits improved resolution and contrast. Gold nanoparticles present in the sample are clearly resolved as dark features.

**Figure 4.**
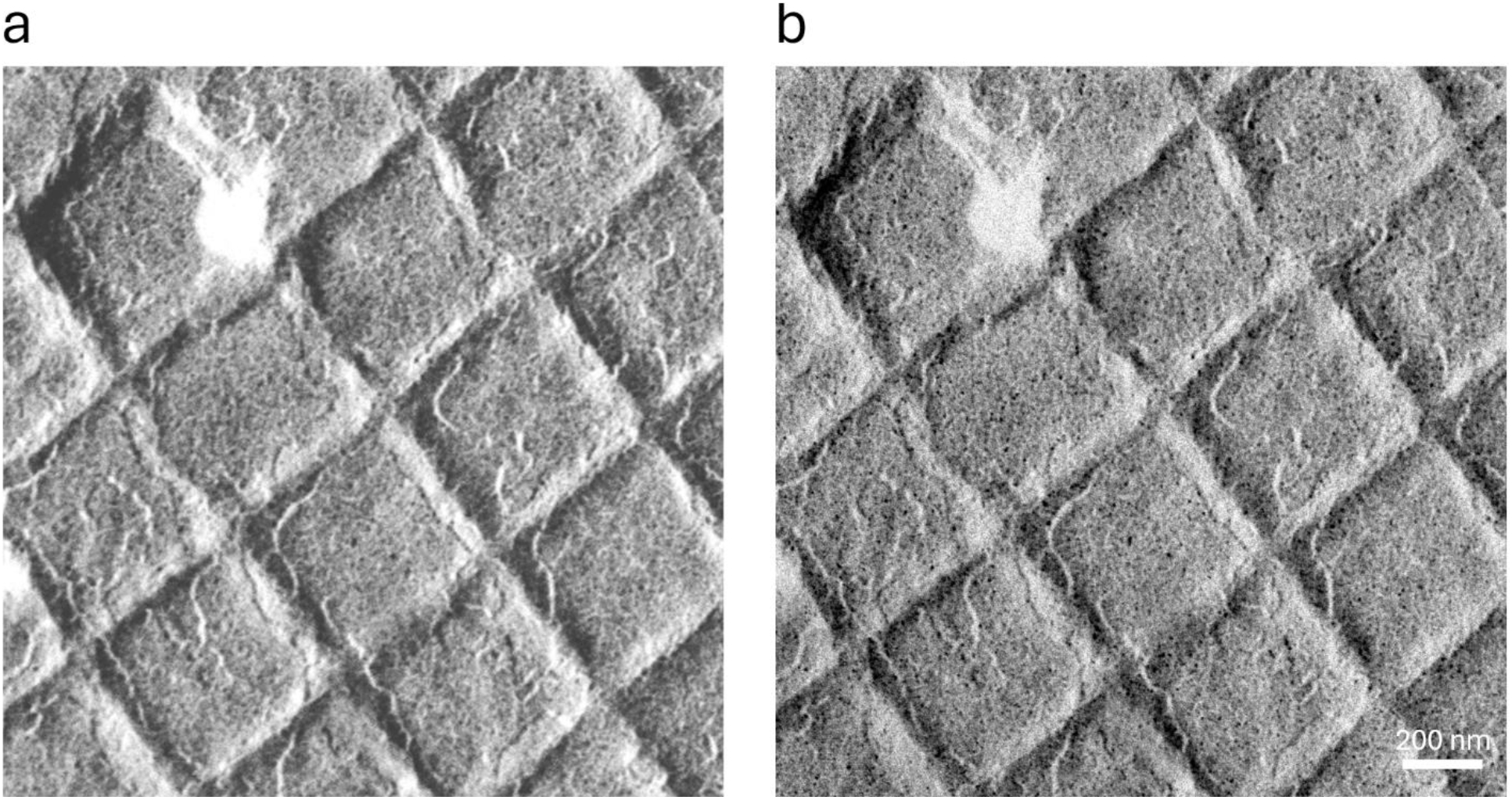
Magnified scan of a replica grating sample with 512 × 512 probe positions. (a) HAADF image, inverted in contrast. (b) Shadow montage based on the 4D-STEM dataset demonstrating upscaled resolution (note the gold nanoparticles as black dots).

The usefulness of the system is demonstrated using a cryo-sample of U2OS cells grown on a gold coated Quantifoil grid. The Tecnai-F20 microscope was set to MicroProbe mode with 70 μm C2 aperture, resulting in a beam semi-convergence of α=1.9 mrad. Using SerialEM with HAADF detector, an overview mapping of the grid in shown in Fig.5a, a square map in Fig.5b, and a scan at the region of interest in Fig.5c. The latter is based on 1024 × 1024 scan points with step size *d*=3 nm/pix. The probe was intentionally overfocused by Δ*f* =10 μm to support formation of shadow images in the diffraction plane. The bright field disc illuminating the pixelated detector covered a diameter of *N*_*BF*_ =53 pixel. A 4D-STEM dataset was acquired for a tilt series in dose symmetric tilt-view order, angles starting at 0, -3, -6, -9, 3, 6, 9, and so forth covering the range between -54 and 54 degrees. The dataset was processed according to the shadow montage technique with synchronization value of *N*_*S*_ =4 according to the formula 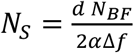[8]. The ShadowMontage_tiltseries_ver2.m script [18] applies CTF flipping to spatial frequencies finer than the probe size, but otherwise preserves the normal bright field contrast, thus providing consistent contrast across scales. The 4D-STEM dataset and resulting aligned shadow montage tilt-series is shared in Zenodo [19]. One tilt view example is shown in Fig.5d, clearly resolving details such as double membranes, cristae, and calcium phosphate granules in three nearby mitochondria [20]. The 3D reconstruction shown in Fig.5e was generated by alignment using IMOD [21] and a subsequent SIRT reconstruction using tomo3d [22]. 3D traces of the cristae, granules, and vesicles are easily observed in the two side projection views.

**Figure 5.**
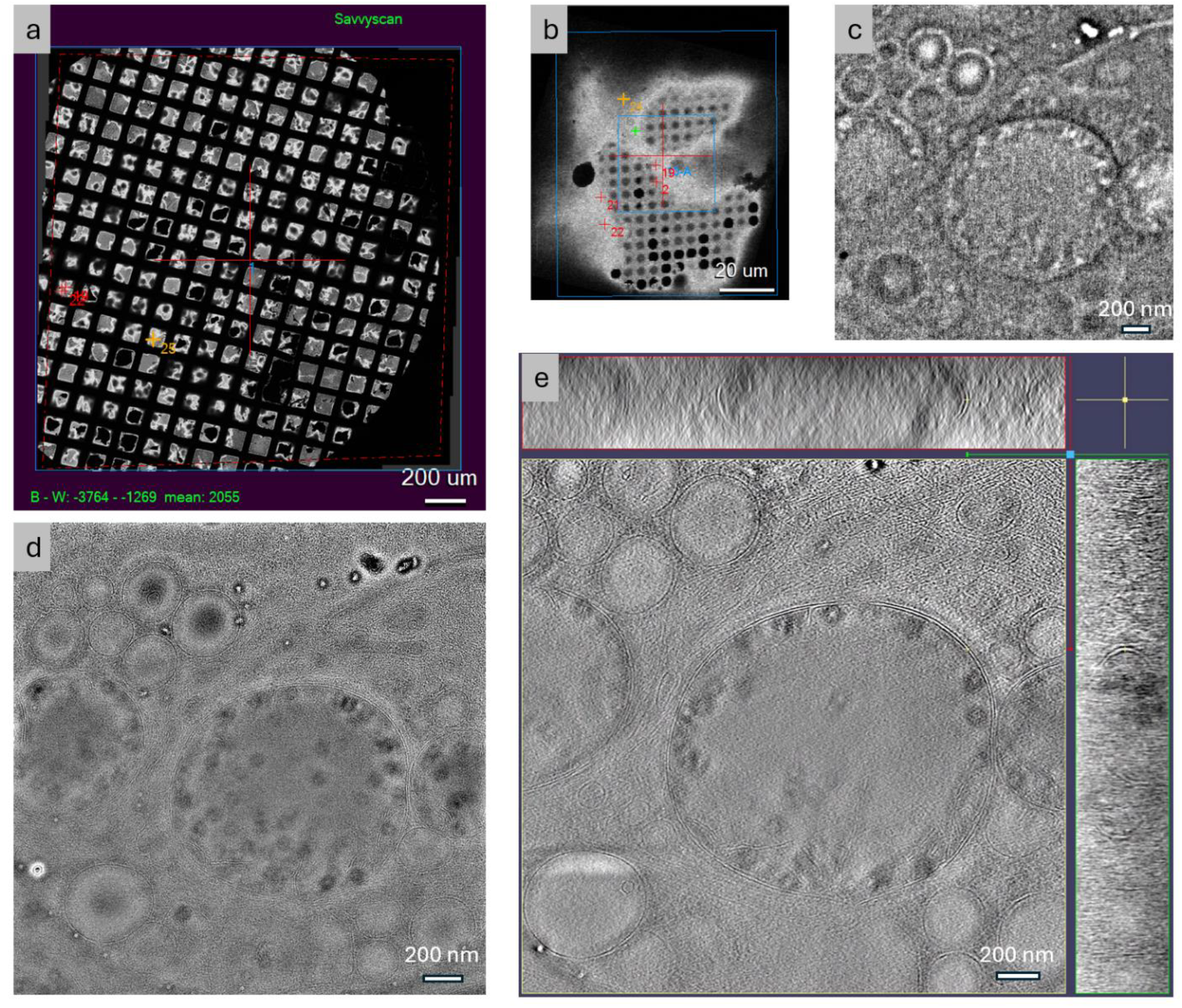
Electron tomography of mitochondria in U2OS cell sample using the hardware integration with SerialEM and a post-processing by shadow montage technique. (a) Overview of the grid generated by SerialEM from a montage of low magnification HAADF scans. (b) One grid-square scan controlled by SerialEM. (c) Direct HAADF image of one projection. (d) Post-processing a shadow montage based on the 4D-STEM dataset of one tilt-view. (e) A tomogram built from 37 tilt-views in shadow montage.

### Conclusions

We have presented a fully open source hardware and code for an FPGA-based system that provides practical acquisition of 4D-STEM data. The hardware is available off the shelf and affordable for any lab, offering further customization for particular needs. Here it was demonstrated for automated tomography. The system and software were designed with SerialEM in mind from the start, so that after installation the operation should be nearly transparent to SerialEM users.

## Appendix

### Technical Implementation

The present system is implemented on a Red Pitaya STEM 125-14 Gen-I board, which is based on a Xilinx Zynq-7010 FPGA. It provides analog-to-digital (ADC) input and dual arbitrary-waveform-generator (AWG) output, both with 14-bit resolution, along with 256 MB of DDR memory. It also includes an integrated Linux development environment, which we use as a server to SerialEM. Adaptations for the Red Pitaya STEM 125-14 Pro Gen-II board, based on the Zynq-7020 FPGA and equipped with 512 MB of memory, are also planned and will be made available in our repository. To drive the scan controller over its full allowable range, the system incorporates a dual-channel JFET operational amplifier (ISL28210S, Intersil). The external trigger output is available on the DIO3_P pin of an extension connector of the board.

Communication between the software stack and the FPGA is handled via several paths. The system bus RAM is used to store control queues and parameters. BRAM is employed for transferring gate patterns, similarly to our previous SavvyGate system. DMA with a FIFO buffer is used to transfer AWG patterns from the software to DDR RAM, and subsequently from DDR RAM to the FPGA. Analog signal acquisition and storage in DDR RAM are based on an FPGA implementation provided by Red Pitaya. Data acquisition is triggered by a pulse on the DIO0_P pin, which can be physically monitored. Further details of the FPGA design are available in the Vivado project that we share on GitHub. All these details are transparent to the user.

### Build Instruction

The instructions below are intended for a specific Red Pitaya board (RPB) STEM 125-14 gen1 connected via a local Ethernet switch to a Windows personal computer (WPC). The internet protocol (IP) name of RPB is denoted by rp-xxxxxx.local.

#### 1. SD card preparation

Prepare the SD card according to the instructions in section 2.5. Prepare SD card — Red Pitaya Documentation with software version (2.07-48).

#### 2. Setting IP addresses

Use a web browser to establish the initial connection. Type rp-xxxxxx.local and select

“Development” and “Web Console” in the graphical user interface. The RPB IP address may be read by command

ip addr

An address that complies with the local microscope network can be set as follows:

nano /etc/systemd/network/20-wired-static-eth0.network

Enter the following content:

[Match]

Name=eth0

[Network]

Address=192.168.100.90/16

Save the file by Ctrl+O and Ctrl+X. Activate the configuration by running

systemctl restart systemd-networkd.service

The IP address and mask of the WPC ethernet socket should be set appropriately. For example, in Ethernet settings change the TCP/IP properties to

IP: 192.168.100.50

Subnet mask: 255.255.255.0

DECTRIS server address should be set accordingly to the content of the start script file IP: 192.168.100.70

Subnet mask: 255.255.255.0

#### 3. SSH connection

To connect with RPB Linux environment, establish SSH authentication between WPC and RPB. Install OpenSSH on the WPC and use PowerShell commands:

ssh-keygen -t rsa -b 4096 (press Enter 3 times).

type $env:USERPROFILE\.ssh\id_rsa.pub | ssh “mkdir -p

∼/.ssh && cat >> ∼/.ssh/authorized_keys” (then answer “root” for password).

The authentication is needed once per WPC. To connect with another PC, repeat the above steps, leaving out the part “mkdir -p ∼/.ssh && “. The command in PowerShell is:

ssh

To copy files from WPC to RPB exit the RPB environment and use:

scp [Windows file]:∼/[Linux folder]/

#### 4. Optional preparation of a new FPGA binary

Download Vivado 2025.2 full version (not lab version), which can be installed with the free Vivado WINPACK license. Download the project from our Github repository and open prj/v0.94/project/redpitaya.xpr. The relevant files are red_pitaya_top.sv, red_pitaya_ps.sv, system.bd and rp_gate_sys_extmem.sv. Double click the system.bd icon in sources explorer to open the system block scheme, which is the hardware component that contains the processing system and our modifications. To compile the changes follow these steps: Validate Design, Generate Output Products, Run Synthesis, Run Implementation, Generate Bitstream. Then copy red_pitaya_top.bif to folder prj\v0.94\project\redpitaya.runs\impl_1, then in the TCL console run

cd C:/AMDDesignTools/RedPitaya-FPGA/prj/v0.94/project/redpitaya.runs/impl_1 bootgen -image red_pitaya_top.bif -arch zynq -process_bitstream bin -w

This will generate a new FPGA binary red_pitaya_top.bit.bin instead of the one we provide.

#### 5. Install the FPGA binary stack

Copy the binary FPGA file red_pitaya_top.bit.bin to the RPB home folder (∼ or /root). On the RPB (Linux), run the following commands from the /root directory:

cp red_pitaya_top.bit.bin /opt/red_pitaya_top/fpga.bit.bin

/opt/redpitaya/sbin/overlay.sh red_pitaya_top x y Full

Check the printed status to confirm that the FPGA is running.

##### 6. Install the C++ SerialEM server software

In RPB Linux make the directories: ∼/SavvyscanPitaya/src, ∼/SavvyscanPitaya/include, and

/opt/velan/Lib. Copy to RPB home folder (∼ or /root) the bash script files compile and start (related to compiling and running the SerialEM server on RPB). Copy the c++ Source files and headers to the Savvyscan src and include folders. Copy the python library content to

/opt/velan/Lib.

Download and install tifffile for the python environment:

pip3 install tifffile-2024.9.20-py3-none-any.whl

Compile the C++ code by running in Linux from /root folder:

bash ./compile

##### 7. SerialEM installation

On a Windows PC intended for SerialEM:

Download and run a 64-bit SerialEM package from SerialEM Downloads and Installation. A subfolder under “Program Files\SerialEM” should be installed.

Allow writing permissions for the SerialEM folder, start a command terminal or PowerShell and cd to SerialEM installation subfolder. Use:

run install.bat (and answer to questions related to your specific microscope). Copy the plugin file Savvyscan.dll to SerialEM folder.

A Windows environment variable SAVVY_SERVER_IP should be set equal to the RPB IP address (In Windows: go to Advanced System Settings, Environment Variables, and add this user variable). The default address is 192.168.100.90, which overrides the address written in SerialEMproperties file.

Create a configuration folder Start_SerialEM which should contain the files SerialEMsettings.txt, SerialEMproperties.txt, SerialEMsettings-scripts.txt, and SEMsystemSettings.txt, which can be copied from our repository. The SerialEMSettings files must be modified in the first lines according to the actual folders, for example: SystemPath C:\Users\seifer\Desktop\Start_SerialEM

ScriptPackagePath C: \Users\seifer\Desktop\Start_SerialEM\SerialEMsettings-scripts.txt

The SerialEMproperties.txt file should be modified according to the actual folders in use and according to the microscope IP address:

SocketServerIP 1 192.168.100.30 #microscope PC address

SocketServerPort 1 48892 #fixed port number

Add a shortcut in folder Start_SerialEM and edit its properties: Target should specify the full path to SerialEM.exe in Program Files and Start in should specify the full path to

Start_SerialEM folder.

## Software Availability

The source code and installation files are available at

https://github.com/Pr4Et/SavvyScan/tree/main/Savvyscan on Red Pitaya

## Acknowledgments

The authors acknowledge support from the Irving and Cherna Moskowitz Center for Bio and Nanobio Imaging, and from the European Union (ERC-Adv CryoSTEM, 101055413; Views and opinions expressed are however those of the authors only and do not necessarily reflect those of the European Union or the European Research Council. Neither the European Union nor the granting authority can be held responsible for them.)

## References

[1] S.G. Wolf, L. Houben, M. Elbaum, Cryo-scanning transmission electron tomography of vitrified cells, Nat Methods 11 (2014) 423–428. 10.1038/nmeth.2842.

[2] D.N. Mastronarde, SerialEM: A Program for Automated Tilt Series Acquisition on Tecnai Microscopes Using Prediction of Specimen Position, Microscopy and Microanalysis 9 (2003) 1182–1183. 10.1017/S1431927603445911.

[3] P. Kirchweger, D. Mullick, S.G. Wolf, M. Elbaum, Visualization of Organelles In Situ by Cryo-STEM Tomography, JoVE (2023) 65052. 10.3791/65052.

[4] S. Seifer, L. Houben, M. Elbaum, Flexible STEM with Simultaneous Phase and Depth Contrast, Microscopy and Microanalysis 27 (2021) 1476–1487. 10.1017/S1431927621012861.

[5] S. Seifer, M. Elbaum, Synchronization of scanning probe and pixelated sensor for image-guided diffraction microscopy, HardwareX 14 (2023). 10.1016/j.ohx.2023.e00431.

[6] P. Kirchweger, S. Seifer, S.G. Wolf, N. Varsano, B. Zens, F.K.M. Schur, M. Elbaum, Azimuthal Segment Imaging in cryo-STEM Tomography, (2025) 2025.11.26.690630. 10.1101/2025.11.26.690630.

[7] S. Seifer, P. Kirchweger, K. Edel, M. Elbaum, Optimizing contrast in automated 4D-STEM cryo-tomography, Microscopy and Microanalysis 30 (2024) 476–488. 10.1093/mam/ozae050.

[8] S. Seifer, L. Houben, Elbaum, Shadow Montage and Cone-Beam Reconstruction in 4D-STEM Tomography, Microscopy and Microanalysis 32 (2026). 10.1093/mam/ozaf126.

[9] B. Küçükoğlu, I. Mohammed, R.C. Guerrero-Ferreira, S.M. Ribet, G. Varnavides, M.L. Leidl, K. Lau, S. Nazarov, A. Myasnikov, M. Kube, J. Radecke, C. Sachse, K. Müller-Caspary, C. Ophus, H. Stahlberg, Low-dose cryo-electron ptychography of proteins at sub-nanometer resolution, Nat Commun 15 (2024) 8062. 10.1038/s41467-024-52403-5.

[10] Y. Yu, K.A. Spoth, M. Colletta, K.X. Nguyen, S.E. Zeltmann, X.S. Zhang, M. Paraan, M. Kopylov, C. Dubbeldam, D. Serwas, H. Siems, D.A. Muller, L.F. Kourkoutis, Dose-efficient cryo-electron microscopy for thick samples using tilt-corrected scanning transmission electron microscopy, Nature Methods 22 (2025) 2138–2148. 10.1038/s41592-025-02834-9.

[11] C. Shi, M.C. Cao, S.M. Rehn, S.-H. Bae, J. Kim, M.R. Jones, D.A. Muller, Y. Han, Uncovering material deformations via machine learning combined with four-dimensional scanning transmission electron microscopy, Npj Comput Mater 8 (2022) 114. 10.1038/s41524-022-00793-9.

[12] B. Kucukoglu, I. Mohammed, R.C. Guerrero-Ferreira, S. Ribet, G. Varnavides, M.L. Leidl, K. Lau, S. Nazarov, A. Myasnikov, C. Sachse, K. Mueller-Caspary, C. Ophus, H. Stahlberg, Low-dose cryo-electron ptychography of proteins at sub-nanometer resolution, (2024) 2024.02.12.579607. 10.1101/2024.02.12.579607.

[13] S. You, G. Varnavides, S. Khavnekar, N. Palatkin, S. Shao, M. Wu, D. Stroppa, D. Chernikova, B. Zhu, R. Egoavil, S. Vespucci, X. Ye, F.K.M. Schur, E. Spiecker, P. Pelz, Gap-free Information Transfer in 4D-STEM via Fusion of Complementary Scattering Channels, (2025). 10.48550/arXiv.2512.19460.

[14] S. Seifer, L. Houben, M. Elbaum, Quantitative atomic cross section analysis by 4D-STEM and EELS, Ultramicroscopy (2024) 113936. 10.1016/j.ultramic.2024.113936.

[15] K. Gong, J. Kim, B.G. Kim, B. Cho, FPGA-based reconfigurable scanning and data acquisition system for scanning electron microscopy, Appl. Microsc. 56 (2026) 16. 10.1186/s42649-026-00137-7.

[16] C.G. Jones, M.W. Martynowycz, J. Hattne, T.J. Fulton, B.M. Stoltz, J.A. Rodriguez, H.M. Nelson, T. Gonen, The CryoEM Method MicroED as a Powerful Tool for Small Molecule Structure Determination, ACS Cent. Sci. 4 (2018) 1587–1592. 10.1021/acscentsci.8b00760.

[17] Nexperion, The SerialEM Script Repository (2026) https://serialemscripts.nexperion.net/category.

[18] S. Seifer, Shadow Montage Scripts, (2026). https://github.com/Pr4Et/Supplementaries/tree/main/Shadow_Montage (accessed June 25, 2026).

[19] S. Seifer, A. Dalaloyan, P. Kirchweger, M. Elbaum, 4D-STEM tilt-series of mitochondria in U2OS cell, (2026). 10.5281/ZENODO.20544863.

[20] S.G. Wolf, Y. Mutsafi, T. Dadosh, T. Ilani, Z. Lansky, B. Horowitz, S. Rubin, M. Elbaum, D. Fass, 3D visualization of mitochondrial solid-phase calcium stores in whole cells, eLife 6 (2017) e29929. 10.7554/eLife.29929.

[21] J.R. Kremer, D.N. Mastronarde, J.R. McIntosh, Computer Visualization of Three-Dimensional Image Data Using IMOD, Journal of Structural Biology 116 (1996) 71–76. 10.1006/jsbi.1996.0013.

[22] J.-I. Agulleiro, J.-J. Fernandez, Tomo3D 2.0 – Exploitation of Advanced Vector eXtensions (AVX) for 3D reconstruction, Journal of Structural Biology 189 (2015) 147–152. 10.1016/j.jsb.2014.11.009.

